# Protein-DNA Clusters Explain the Non-Exponential DNA Residence Time Distributions of Transcription Factors

**DOI:** 10.1101/2023.09.21.558872

**Authors:** Zafer Kosar, Aykut Erbas

## Abstract

The transcription process is regulated by temporal interactions of transcription factors with DNA. In the last decade, computational and experimental studies revealed the residence times of transcription factors on DNA correlate with transcriptional output. Biochemical studies suggest that transcription factor bindings exhibit bi-exponential dynamics, often explained by the binary affinity model composed of nonspecific and specific protein-DNA interactions. Recently, transcription factor residence times were shown to display a power law *in vivo* implicating effective protein-DNA interactions controlling the dissociation kinetics are rather more complex than suggested. One contribution that can cause such continuous residence-time distributions could be higher-order protein-DNA complexes or protein coacervates. Here, by using molecular dynamics simulations of a coarse-grained polymer model for bacterial chromosomes interacting with homodimeric transcription factors at physiologically relevant concentrations, we demonstrate that residence time distributions of dimeric proteins follow a multi-exponential pattern even when a single interaction describes the affinity between DNA and protein. Our simulations reveal that this emergent behavior is due to the formation of DNA-protein clusters of various sizes at a wide range of protein concentrations and affinities. These findings add another layer to transcriptional regulation and, consequently, to gene expression by connecting transcription factor concentrations and affinities, DNA-protein clusters, and DNA residence times of transcription factors.

## Introduction

Many aspects of cellular behavior, including cellular identity and response to physiochemical and even biomechanical stimuli, are dictated by gene expression patterns. Among the many regulators of gene expression, transcription factors (TFs) play one of the most prominent roles in inhibiting or activating the transcription of target genes at the single-binding site level (Browning, 2004; Seshasayee et al., 2011; Bintu et al., 2005; Gentry et al., 2023). TFs locate their specific DNA binding sequences through random search. While 3-dimensional (3D) diffusion and intersegmental transfer of TFs are possible (Halford and Marko, 2004), the search process is driven mostly by 1-dimensional (1D) sliding along nonspecific DNA sites, suggesting that TFs remain DNA-bound for most of their lifetimes (Stracy et al., 2021).

Regulation of a gene by a TF starts at the binding site once the TF binds to its specific DNA sequence and forms a relatively stable protein-DNA complex. This complex controls the repression or activation of the gene until the protein unbinds from the DNA binding site via thermal fluctuations or collision with other molecular species (de Jonge et al., 2022; Haberle and Stark, 2018). Accordingly, recent studies in TF dynamics revealed that the residence time of a TF on its DNA binding site (i.e., its unbinding or dissociation rate with the units of inverse time) is directly intertwined with transcriptional output (Clauß et al., 2017; Lickwar et al., 2012; de Jonge et al., 2020). Thus, the factors affecting the DNA-residence time of TFs emerge as key components in the gene expression patterns.

The duration of a TF on its DNA binding site can depend on various factors, including TF’s affinity to the DNA (Connaghan-Jones et al., 2007; Schaaf and Cidlowski, 2003), temperature (Bonner et al., 1992; Zhu et al., 2023), and 3D DNA structure (Kim and Shendure, 2019; Inukai et al., 2017). Studies focusing on unbinding (or dissociation) kinetics of TFs from their single-binding sites have demonstrated a strong concentration dependency of TF unbinding rates Graham et al. (2011); Joshi et al. (2012); Kamar et al. (2017). In scarce concentrations, a TF remains bound to its target site for extended periods. In contrast, the abundance of unbound TFs in solution leads to a competition for the binding sites on DNA, resulting in much higher unbinding rates (i.e., shorter residence times) via the process referred to as facilitated dissociation (FD) (Kamar et al., 2017; Koşar et al., 2022; Erbaş and Marko, 2019; Joshi et al., 2012; Graham et al., 2011; Dahlke and Sing, 2017).

The impact of TF concentration is not limited to protein unbinding dynamics via FD. TFs can also contribute to 3-dimensional (3D) genome organization and form DNA-protein clusters in a concentration-dependent manner (Kim and Shendure, 2019; Noort et al., 2004; Skoko et al., 2006, 2004; Remesh et al., 2020; Koşar and Erbaş, 2022; Arold et al., 2010; Dame et al., 2000; Winardhi et al., 2015). Such structural effects are more pronounced with bacterial Nucleoid-Associated Proteins (NAPs), a class of DNA-binding proteins often with dual functionality (i.e., regulatory and structural roles). NAPs are involved in chromosome organization, many of which also function as transcription factors (Dillon and Dorman, 2010; Dorman, 2014). Due to their common multivalent nature (i.e., multiple DNA binding domains) Lee (1992), DNA-binding proteins can drive the bridging of distinct DNA segments and the formation of DNA-protein clusters of various shapes and sizes and other forms of chromosome architectural changes (Skoko et al., 2004; Dillon and Dorman, 2010; Wang et al., 2011; Hammel et al., 2016; Dame, 2005; Verma et al., 2019). Molecular Dynamics (MD) simulations of model bacterial systems suggest that these events are highly dependent on protein concentration and nonspecific interaction potential (i.e., the affinity between the DNA-binding protein and nonspecific DNA sites) (Koşar et al., 2022; Brackley et al., 2013; Jung et al., 2024; Dahlke and Sing, 2019).

Recent experimental studies employing single molecule tracking (SMT) demonstrated that the residence duration of several TFs and chromatin-associated proteins dynamics follow a power-law pattern (Garcia et al., 2021; Garcia, 2021). These findings support a continuum model for TF dynamics and TF-DNA interaction affinities rather than a bi-exponential model attributed to binary specific (*U*_*sp*_) and nonspecific (*U*_*ns*_) affinities. In the bi-exponential model, TFs are considered to have longer residence times on their specific target DNA sequence and shorter residence times on nonspecific DNA, often resulting in the utilization of a bi-exponential decay equation to explain the distribution of residence times (Chen et al., 2014; Ball et al., 2016; Morisaki et al., 2014). Contrary to this suggestion, even single affinity (*U*_*ns*_ = *U*_*sp*_) and binary affinity models (*U*_*ns*_ *< U*_*sp*_), in principle, can generate a non-trivial dissociation energy landscape via many-body interactions via formation of protein-rich phases through liquid-liquid phase separation (Boija et al., 2018; Hnisz et al., 2017; Sabari et al., 2018; Wang et al., 2021; Nozawa et al., 2020; Larson et al., 2017; Alberti et al., 2019).

Here, we employed a topologically accurate large DNA polymer and state-of-the-art protein models with varying levels TF-DNA affinity and protein concentrations for extensive MD simulations. Analysis of our simulations revealed apparent multi-exponential distribution patterns even with the single and binary affinity models, hinting at emergent behavior. Consequently, we explored the driving factors of multi-exponential residence behaviors of TFs. In particular, high nonspecific affinity cases, which enabled the formation of much larger clusters, exhibited distinct RT distributions and required more exponents to be matched with decay curves. Investigation of TF-DNA cluster formations of different sizes revealed that cluster dissipation means lifetimes are coupled to their sizes.

Therefore, concentration and affinity-dependent cluster formations could be the driving factors of the observed multi-exponential patterns.

We then explored the distributed affinity models with uniform or normal distributions to check whether the additional complexity via mimicking the continuum model would lead to power-law behavior. However, it is clear that discrete affinity distributions with limited resolutions again form multi-exponential decay patterns, and thus, they do not provide sufficient complexity to generate a power-law behavior.

In this study, we demonstrate that TF dynamics follow multi-exponential patterns even without multiple TF-DNA affinity levels in our model. In parallel with our findings, multi-exponential fits were previously used in several SMT studies to interpret the residence behaviors (Hipp et al., 2019; Reisser et al., 2020; Agarwal et al., 2017). We also show how TF residence patterns are impacted by DNA-protein cluster formations. Our model predicts the power-law behavior might be plausible even with discrete affinities with some additional complexities or with higher resolution of affinities. These findings may help establish a more advanced understanding of transcriptional regulation and, thus, of gene expression regulation.

## Results

### Residence time behaviors are multi-factorial

We employed multiple cases for nonspecific interaction affinities ranging from a very weak 1*kT* to a strong 4*kT* (i.e., *U*_*ns*_ = *U*_*sp*_) per bead, where *U*_*sp*_ = 4*kT* per bead in all cases. For each of these affinity levels, four different physiologically relevant (Verma et al., 2019; Azam et al., 1999; Ball et al., 1992) concentrations ranging from 10 *−* 60*μM* of TFs were employed in the simulations. We obtained residence time patterns, as shown in Figure 1 and as described in methods, in the form of occurrence versus duration, where occurrence is the number of times a duration was achieved or observed. We then tried fitting several equations to define TF residence patterns (Figure 1E). Fit equations included exponential decay with up to five exponents (Figure 2). The projection of RT patterns in the log-log scale exhibited an apparent arching pattern, eliminating the possibility of a good power-law fit, which produces a straightline pattern in that scale. The single exponential decay (ED) equation can only describe short-duration (*<* 10*a*.*u*.) distributions regardless of affinity and concentration (Figure S1), implying the full-spectrum residence time behavior is not dependent on a single parameter. This behavior was indeed expected due to distinct non-specific and specific affinity levels leading to at least a bi-exponential behavior.

**Figure 1.**
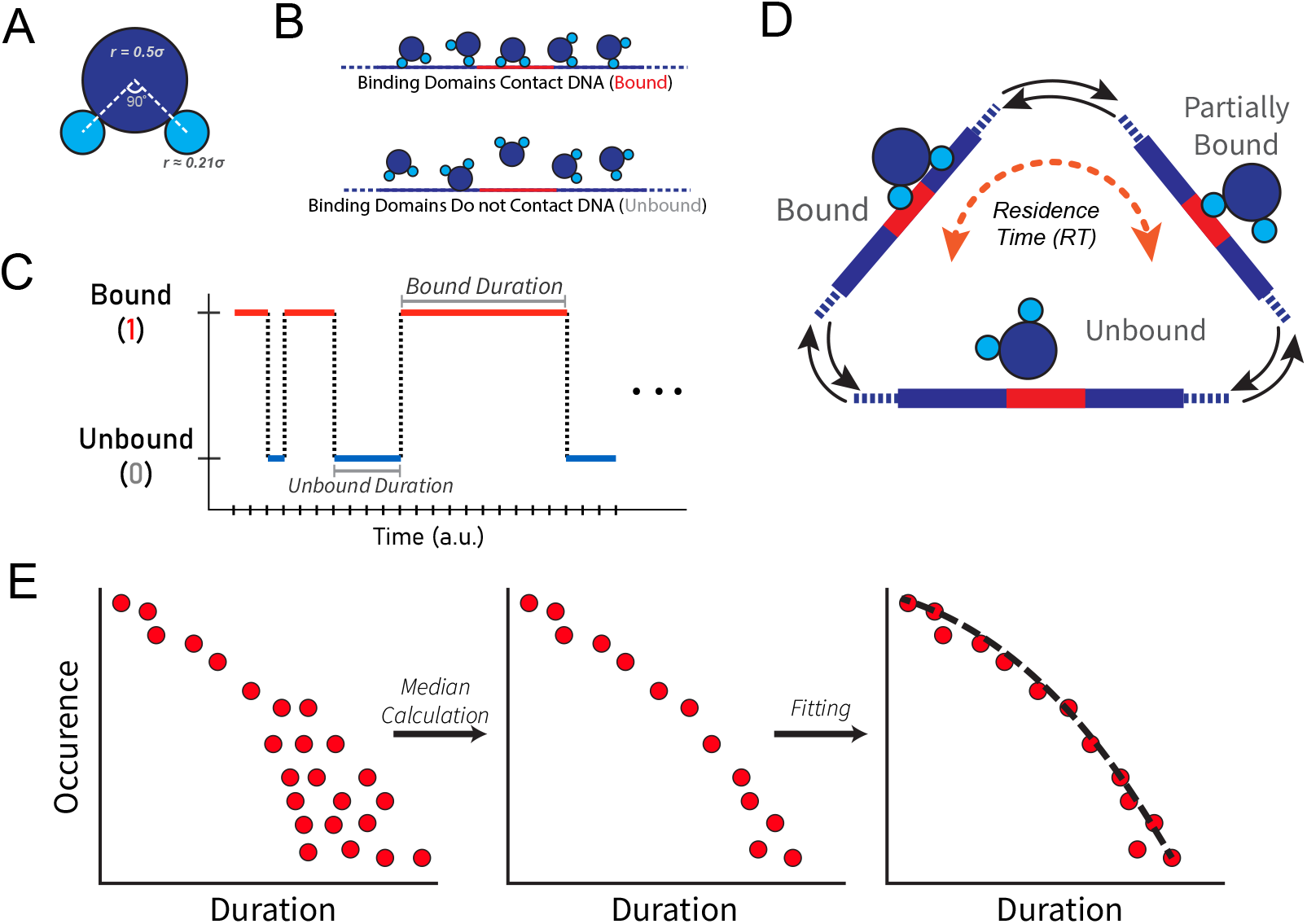
Graphical abstract of the study. (A) The coarse-grained model of a generic homodimeric transcription factor. The light blue parts represent the binding domains and the dark blue part is the hinge (non-binding) domain. Radii of the coarse-grained beads are given in Lennard-Jones (LJ) units and the angle between binding domains are shown in degrees. (B) Distinction between bound and unbound states. (C) Graphical representation of DNA-residence durations. (D) Possible binding states of a transcription factor and calculation of residence times (RT). A transcription factor could be fully or partially bound to DNA, or it could be in an unbound state in which it does not interact with the DNA. Note that red and dark blue regions represent specific binding sites and nonspecific binding sites of the DNA, respectively. Nevertheless, binding to either site is considered equivalent. (E) Collection, median calculation, and analysis of transcription factor residence patterns. Bound durations of each transcription factor throughout the simulations are collected and then minimized to their medians. Resulting patterns are analyzed by fitting several exponential equations.

**Figure 2.**
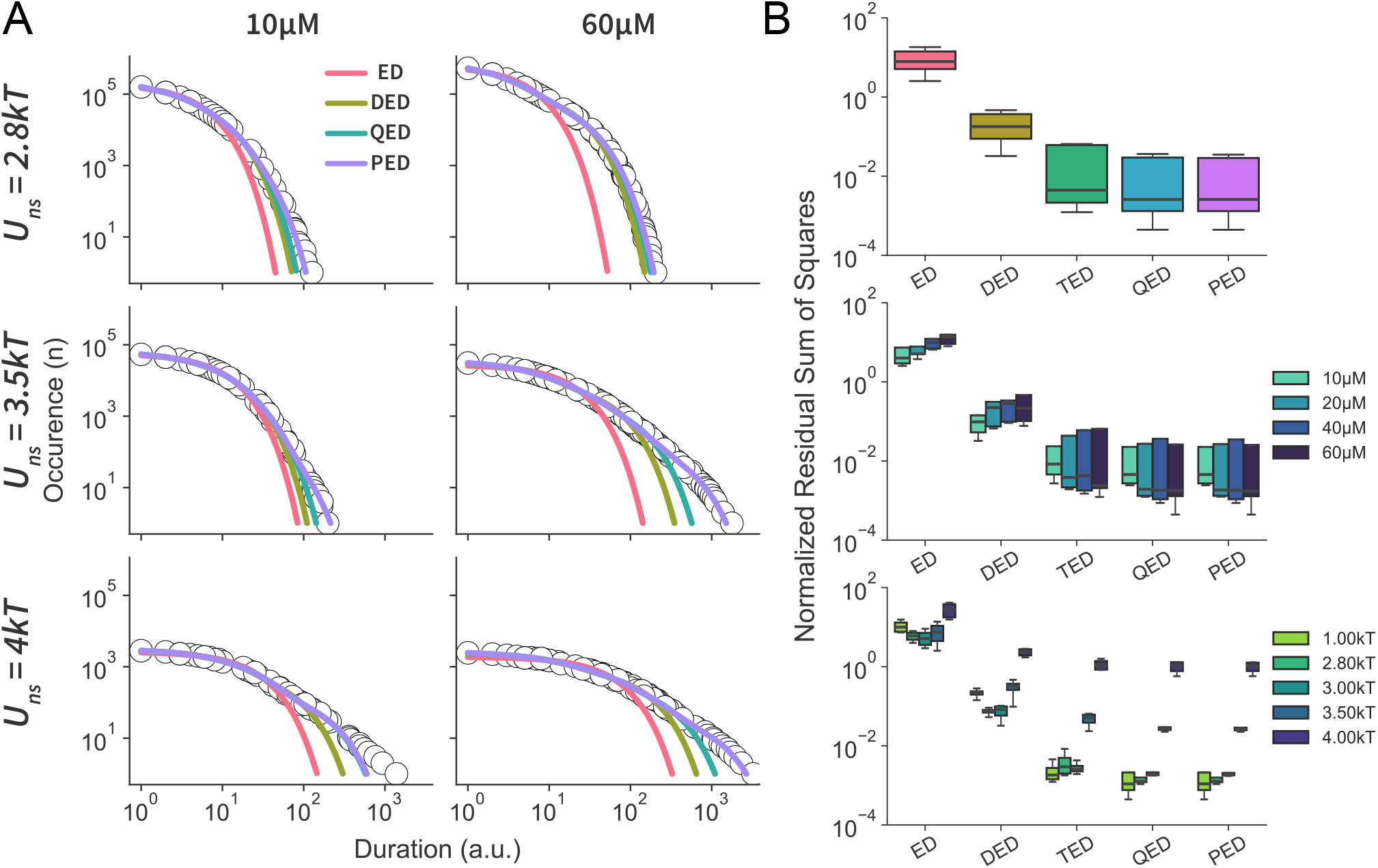
Analysis of transcription factor residence time patterns. (A) Distribution of transcription factor residence times in arbitrary unit (a.u.) at 10*μM* and 60*μM* concentrations for distinct *U*_*ns*_ levels and the fits of Exponential Decay (ED), Double Exponential Decay (DED), Quadruple Exponential Decay (QED), and Pentuple Exponential Decay (PED) equations to residence time data. Due to the extensive number of data points, only representative 20% are shown here. (B) Normalized Residuals Sum of Squares (RSS) of the fits depicting the scale of difference between observation and the fits. For all equations, overall RSS, RSS by concentrations, and by binding affinities are shown, respectively. Note that specific binding energies are 4*kT* for all the systems.

For most of the cases, our simulations utilized binary values where, for each simulation, there was only one nonspecific and one specific affinity where *U*_*ns*_ *< U*_*sp*_. If there was no emergent factor affecting residence times, a double exponential decay (DED) would be sufficient to interpret the residence time patterns obtained from the simulations. Respectively, we used a double exponential decay equation for characterizing RT patterns. DED provided much better fits compared to the ED equation but notably failed to generalize well for all durations (Figure 2A), which is intriguing considering simulations utilized a single type of TF and two distinct types of DNA sites.

The strategy employed here was to increase the number of exponents to better characterize TF residence behaviors. We gradually increased the number of exponents for the decay equation to up to five exponents. On top of the visual inspection of the fits, we quantitatively analyzed their accuracy via normalized Residual Sum of Squares (RSS), where the lower RSS indicates a better fit. The increase from single to double as well as from double to triple exponential decay resulted in the order of magnitudes lower RSS (Figure 2B). Additional increments in the number of decay exponents also reduced RSS for the fits, but the changes were not as drastic. These analyses demonstrate that RTs follow a multi-exponential decay pattern, suggesting RT patterns are shaped by additional factors besides binary DNA-protein binding affinities.

### Residence time behaviors are dependent on the emergent behavior of TF concentration and binding energies

Residence time behaviors (Figure 2A, Figure S1) suggest a pattern beyond a bi-exponential system that cannot be simply explained nor attributed to the dual model of short-lived TF-DNA interactions on nonspecific sites and longer-lived, more stable interactions on specific binding sites. Therefore, an emergent behavior is required to explain such multi-exponential patterns. Specifically, patterns obtained from elevated concentration and high-affinity (*kT*) cases require more exponents (Figure2A, Figure S1), indicating such a behavior emerges as a result of concentration and energy levels. The relatively high binding energies and concentration of DNA-binding proteins were shown to lead to cluster formations as well as local and global condensations of the chromosome (Koşar et al., 2022; Lin et al., 2012; Brackley et al., 2013; Agback et al., 1998). Unsurprisingly, higher concentrations yield larger clusters (Figure 3C,E). Remarkably, global compaction of the chromosome and cluster sizes, even conformations, are mainly regulated by nonspecific interactions, and specific interactions have little effect (Koşar et al., 2022). This behavior can be attributed to the abundance of nonspecific DNA sites over specific sites. In this work, we reduced the impact of global chromosome compaction via miniaturizing TF binding domains (Figure 1A), which minimizes the bridging of multiple DNA segments. Moreover, the use of adequate binding energies eliminated the possibility of high chromosomal compaction even at high concentrations (Figure 3A). This strategy allowed TFs to roam relatively freely within the cellular confinement.

**Figure 3.**
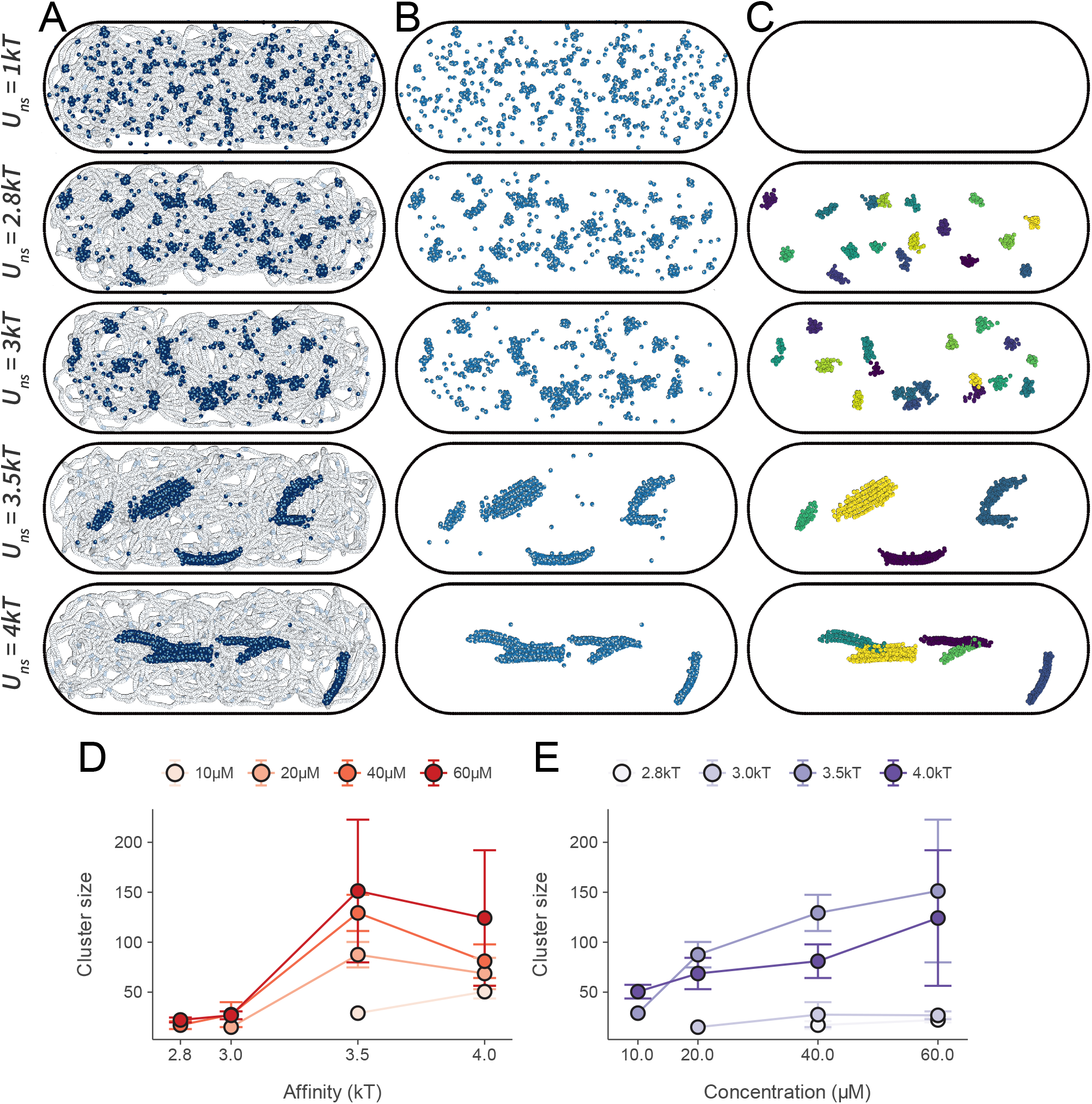
Visualizations of the system, protein distributions, and cluster formations of the coarse-grained bacterium model at various nonspecific binding energies. (A) Overview of the system DNA (white-light blue), and transcription factors. (B) Transcription factor distributions within the confinement. (C) Cluster formations of transcription factors. Here, the coloring only serves to distinguish distinct protein clusters. Snapshots were obtained from systems with transcription factor concentrations of 60*μM*. Specific binding energies are 4*kT* for all the systems. (D) Distribution of cluster sizes (number of transcription factors) by nonspecific TF-DNA affinities or (E) by TF concentrations.

Our model depicts TFs residing for increasingly higher durations on DNA instead of freely roaming around with the increasing nonspecific potential with a fixed specific binding energy (Figure 3A,B). Consistent with the previous findings, at a very low nonspecific potential (*U*_*ns*_ = 1*kT*), bound proteins are sparsely distributed around the DNA polymer, mainly near binding sites (Figure 3A), but do not form DNA-polymer complexes (i.e., clusters) (Figure 3C). Relatively higher nonspecific potentials (*U*_*ns*_ = 2.8*kT* and *U*_*ns*_ = 3*kT*) led to primarily small and globular clusters. In contrast, stronger non-specific affinities (*U*_*ns*_ = 3.5*kT* and *U*_*ns*_ = 4*kT*) enabled the formation of much larger clusters (Figure 3D) with filamentous conformation (Figure 3C). Therefore, we hypothesized the cluster sizes and conformations to play significant roles in TF residence patterns.

Cluster formations drive multi-exponential residence time patterns

To unravel the relationship between DNA-protein clusters and multi-exponent patterns, we initially considered tracking and collecting residence times of TFs for each individual cluster, as described in Figure 1. However, clusters are dynamic formations, and it is not rational to follow TFs that were initially part of a cluster because after they dissipate, they are free to bind anywhere on DNA. Therefore, we rather investigated the dissipation mean lifetimes of individual clusters (see methods). As opposed to RT pattern acquisitions, partial unbinding events were also counted. This modification was needed simply because at high nonspecific affinities, the time needed for obtaining cluster decay rates would exceed simulation lifetimes when only full unbinding events were counted.

At relatively low nonspecific energies (*U*_*ns*_ = 2.8*kT* and *U*_*ns*_ = 3*kT*), correlation analysis of cluster size and their dissipation durations did not reveal any relation. Contrarily, higher nonspecific interaction potentials (*U*_*ns*_ = 3.5*kT* and *U*_*ns*_ = 4*kT*) led to correlations with a coefficient of *r ≥* 0.5 (Figure 4), where larger clusters had longer mean lifetimes. These imply differently sized clusters could dissipate at diverse rates, contributing to multi-exponential behaviors that are more apparent for high nonspecific affinity cases (Figure 2A).

**Figure 4.**
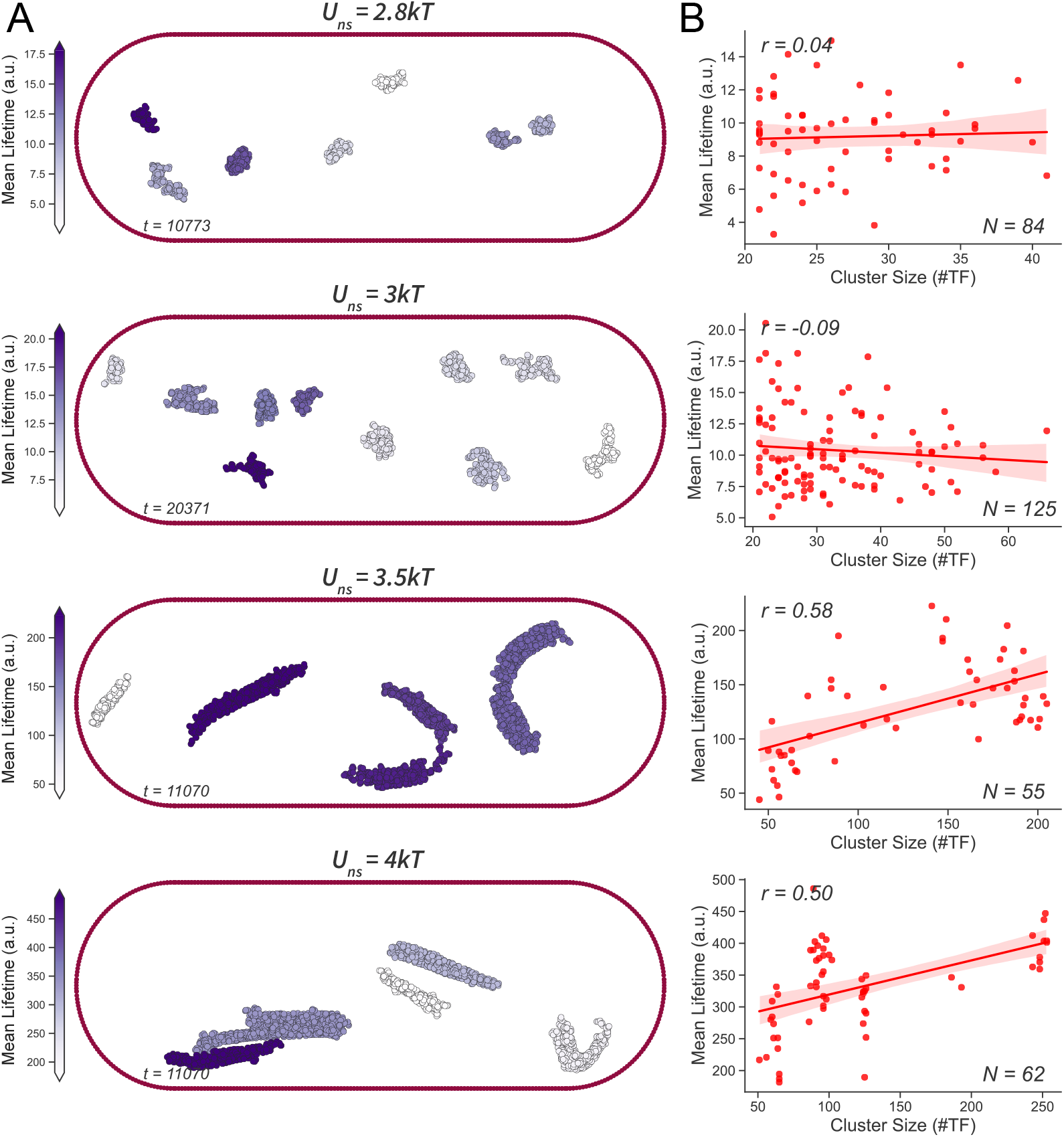
Correlation analysis of cluster sizes and dissipation times. (A) Snapshots of the transcription factor clusters at given nonspecific energies from random timesteps. Darker colors implicate a higher mean lifetime, as indicated by the color bars on the left. (B) Mean lifetimes in arbitrary units versus cluster sizes in the number of transcription factors they are composed of. Regression lines are used to determine correlations. Pearson correlation coefficients (r) are given on the upper-left side, and a number of the clusters in the analysis is given on the bottom-right side. Semi-transparent bands around the regression lines depict 95% confidence intervals. Transcription factor concentrations were 60*μM* for all cases. The threshold was set to a minimum of 20 TFs for cluster size for decay rate analysis for a more reliable statistical approach. 12 time points were used for sampling and the specific binding energy was 4*kT* for all simulations.

It should also be noted that at 2.8*kT* and 3*kT* nonspecific affinities, cluster sizes ranged from 20 to 45 and 20 to 70 TFs, respectively (Figure 4B). The sizes were significantly improved with higher nonspecific interactions. The cases of 3.5*kT* and 4*kT* nonspecific affinities allowed the formation of clusters of size ranging up to 250 TFs (Figure 4B). This increase, of course, provides a statistical advantage for the analysis of the relation between lifetimes and cluster sizes.

### TF residence times depend on their position relative to clusters

Upon establishing the relationship between cluster sizes and their lifetimes, we considered analyzing the residence times of TFs that are located at different regions of the clusters. Cluster-associated TFs can be classified as surface TFs and core TFs (Figure 5B). While the core TFs are rather trapped within the cluster, surface TFs are more exposed and could be expected to have shorter residence durations as they are free to unbind. Additionally, TFs that are not part of any cluster are classified as free TFs (Figure 5B), which are useful as references against cluster-associated TFs.

**Figure 5.**
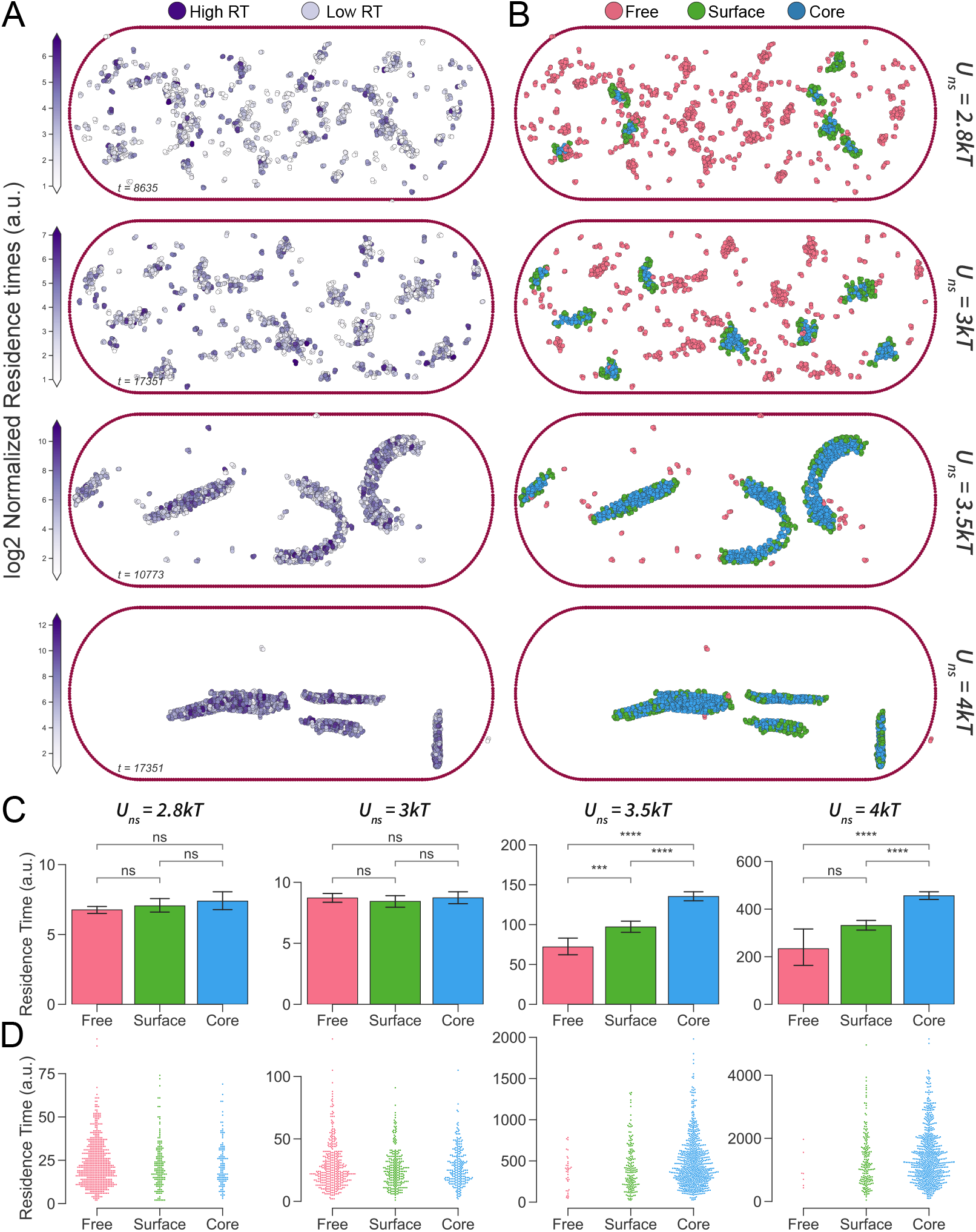
Residence time analysis based on the transcription factor localization with respect to clusters. (A) log2 normalized residence times of individual transcription factors at various nonspecific DNA-TF interaction affinities at 60*μM* TF concentration (B) Classification of TFs according to their positions with respect to the clusters. ‘Free’ transcription factors do not belong to any clusters. Transcription factors on the ‘Surface’ and in the ‘Core’ of the clusters were named accordingly. (C) Differences among residence times for free, surface, and core transcription factor types at given nonspecific affinities. Error bars represent 95% confidence intervals. Welch’s *t-test* was used for significance analysis. (D) Distributions of TF residence times for the given transcription factor types at 60*μM* cases. 12 random timesteps were used for sampling, and the specific binding energy of 4*kT* was used for all simulations. Representative 12% of the data is shown to prevent over-crowding in swarm plots.

Visual investigation of individual proteins located at different positions with respect to the TF-DNA clusters did not show apparent distinction in their residence times (Figure 5A). However, in an analytic (i.e., Welch’s *t-test*) and more comprehensive approach utilizing multiple time points, core TFs exhibited significantly longer residence times compared to surface and free TFs (Figure 5C). However, this behavior was limited to the high nonspecific affinity cases and was not observed at lower nonspecific potentials, which may add another layer of complexity to residence behaviors and, therefore, could explain the more prominent multi-exponential residence patterns at such energy levels. At *U*_*ns*_ = 3.5*kT* case, surface TFs also had significantly higher residence times in comparison to free TFs (Figure 5C).

This disparity indeed reveals that even located at the exposed regions of clusters, TFs could behave differently compared to freely roaming TFs. Thus, the variations among core, surface, and free TFs residence times could shape the overall residence distributions (Figure 5C-D) and further contribute to multi-exponential DNA-residence behaviors of TFs at single and binary affinity models.

Another factor that should be considered is the differing number of TFs of each type for distinct cases. Even though the concentration (60*μM*) and specific binding potentials (*U*_*sp*_ = 4*kT*) are fixed among the cases, the higher nonspecific energies drive clustering rather than scattering (Figure 3B), also leading to the formation of much bigger clusters (Figure 3C-E, Figure 4). Thus, more of the TFs are located within the clusters for high nonspecific affinity cases (Figure 5D). Moreover, the larger the clusters, the higher the core TF ratios were (Figure 5D), as expected due to the decrease in the surface-to-volume ratio.

### Distributed affinity models do not lead to power-law behavior

Our MD simulations revealed that even single and binary affinity models could lead to multi-exponential residence time patterns. The next step was to mimic the continuum affinity model with a distributed affinity model, where we employed 13 distinct DNA site types and assigned them a range of DNA binding affinities. These affinities inclusively ranged between 1 *−* 4*kT* and 2 *−* 5*kT* with an increment of 0.25*kT* and the number of DNA sites for the given affinity assigned to provide either a normal (i.e., Gaussian) or uniform distribution.

Contrary to our expectations, distributed affinity models did not produce power-law behavior. Instead, the triple-exponential fit, an example of multi-exponential fit, better described the behaviors (Figure 6). Therefore,we might speculate that mimicking the continuum model via discrete affinity distribution models is not sufficient for power-law behavior, and such behavior would require much higher complexity and resolution than our current systems for MD simulations could provide. On the other hand, the extraordinary complexity of biological systems or continuous affinity levels with higher resolutions (higher number of distinct affinity levels) could be sufficient to generate such behavior and might be the underlying reason behind the power-law behavior in the experimental approaches.

**Figure 6.**
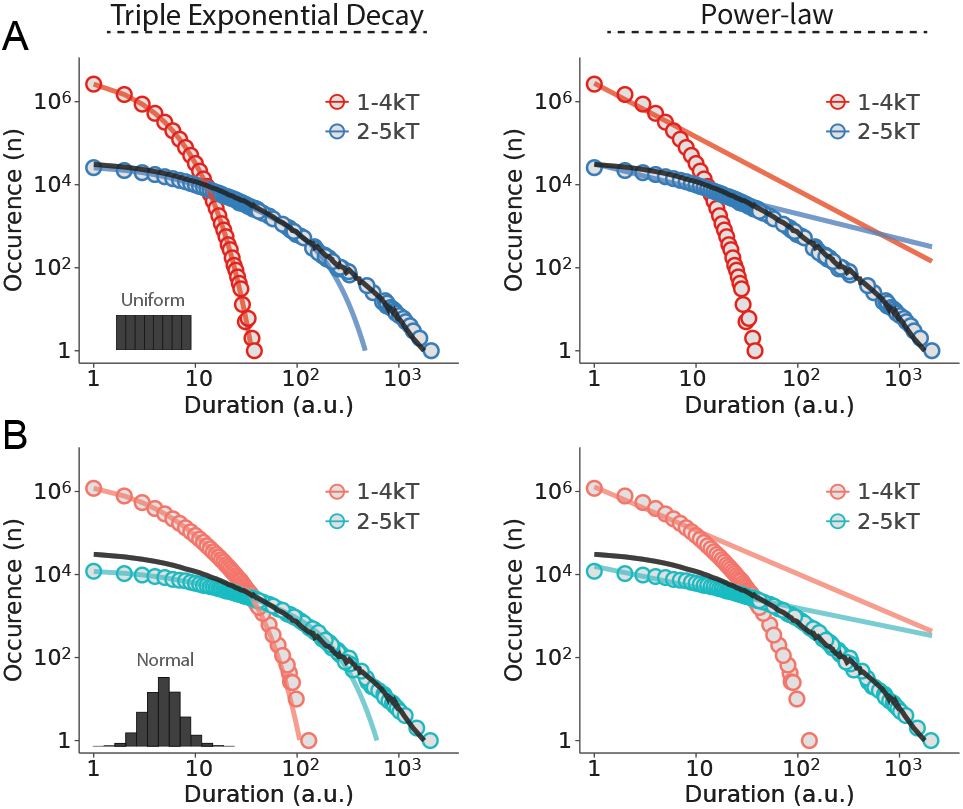
Transcription factor residence time behaviors at distributed affinities. (A) Uniform distribution and (B) Normal (Gaussian) distribution affinity models. Pattern analysis via fitting of triple exponential decay (left column) and powerlaw (right column) respectively. Example histograms of the uniform and the Gaussian distribution are shown in the bottom-left corner of each panel’s first graph. Transcription factor concentrations were 60*μM* for all cases. Black lines are shown as a reference from the nonspecific affinity of 3.5*kT*.

## Discussion

The durations, therefore the distributions of durations of TFs on DNA have a significant impact on transcriptional regulation (Clauß et al., 2017). Using extensive MD simulations with a large-scale DNA polymer and state-of-the-art transcription factor coarse-grained models enclosed within rod-like confinement, we investigated the residence time distributions of the homodimeric generic TFs. In our investigations, we employed several biologically relevant TF concentrations (Ishihama et al., 2014) and distinct binding strengths between TFs and DNA. Our findings reveal an apparent multi-exponential behavior for all of the single, binary, and distributed affinity models of TF-DNA bindings. Single and binary affinities leading to multi-exponential patterns were beyond the prior suggestion of bi-exponential behavior due to specific and nonspecific binding sites. Here, we demonstrate how these multi-exponential behaviors are shaped by TF-DNA complexes, reveal the driving factors of such formations, and connect the residence times behavior to nonspecific binding affinities.

Within the context of their primary roles, TFs can inhibit and activate transcription. The formation of stable TF-DNA complexes extends the duration of a TF’s residence time on its target DNA site. That, in turn, could significantly enhance their inhibitory or activatory effects. We may also speculate that larger and more compact clusters may decrease the accessibility by RNA polymerase, such as in heterochromatin-like domains (Amemiya et al., 2022), effectively reducing the transcriptional output. Contrarily, similar protein-DNA complexes, but in the form of transcriptional machinery, could initiate or enhance transcription.

The affinity of a transcription factor to its specific binding site is often stronger compared to its binding affinity towards a nonspecific DNA sequence. However, nonspecific DNA-TF interactions can contribute to the global chromosome organization due to their abundance. Collectively, specific and nonspecific interactions can lead to the formation of local DNA-protein complexes (i.e., clusters) (Koşar et al., 2022; Lin et al., 2012; Brackley et al., 2013; Agback et al., 1998). Moreover, these nucleic acid-protein complexes could significantly affect transcriptional regulation. For instance, transcription factories are multiprotein complexes formed by TFs, RNA polymerase, coactivators, etc. (Iborra et al., 1996; Cook, 2010; Melnik et al., 2011; Mitchell and Fraser, 2008). Therefore, multiprotein complexes or clusters are crucial regulators of transcriptional output. Furthermore, eukaryotic euchromatin regions, which are not densely packed as heterochromatin regions, are more accessible to transcription factors and thus more transcriptionally active (Amemiya et al., 2022; Elgin and Grewal, 2003; Penagos-Puig and Furlan-Magaril, 2020), emphasizing the significance of local DNA organization on residence times. In other words, the contribution of DNA-binding proteins to gene regulation is not restricted to their primary functions as TFs; their activity in domain-specific genome organization is also crucial.

Even though we demonstrated cluster size is an important factor for residence times, we also wanted to reveal any possible relationship between cluster shape (conformation) and dissipation rates. However, investigation of the conformations of the cluster formations on residence times was inconclusive due to cluster sizes being highly intertwined with nonspecific affinity levels. Thus, establishing a clear relation between cluster shape and RT requires eliminating affinity as a variable. In turn, that would require a separate MD model. Furthermore, clusters are not fixed formations; they are highly dynamic. Their sizes, conformations, orientations, and even the DNA segments they interact with change over time in the system. Moreover, clusters can merge and split, making it rather complicated to track individual clusters and make accurate analyses (Wang et al., 2011).

The utilization of dimeric TF models might have contributed to the multi-exponential behavior in two ways. First, our definition of unbound TF requires both binding domains not to be in touch with any DNA molecule (Figure 1B). In other words, partial bindings to DNA (i.e., bindings with only one binding domain) are equivalent to full bindings in terms of being considered bound. The difference in stability of partial binding and full binding may indeed lead to different residence durations contributing to multi-exponential behavior. Moreover, divalent interactions are significantly more prone to facilitated dissociation (Kamar et al., 2017; Chen et al., 2018). Since FD is pronounced at high concentrations (Koşar et al., 2022), multi-exponential behavior could be partially attributed to FD as it can facilitate the unbinding of the exposed TFs (i.e., free or surface) more than the unexposed TFs (i.e., core).

Noise in gene expression is an important factor driving cellular heterogeneity (Liu et al., 2019). Gene expression noise could lead different cells in a homogeneous population to distinct phenotypes even under the same environmental conditions (Raser and O’Shea, 2005). This resulting cellular heterogeneity may provide evolutionary advantages to cells. Notably, the primary driving event of gene expression noise is considered to be TF binding (Parab et al., 2022). Also, infrequent or rare biochemical processes contribute to noise in gene expression (Raser and O’Shea, 2005). Such infrequency or lower occurrence could be seen for the higher duration residences in our systems, which are more apparent at high nonspecific affinity and high protein concentration cases. Therefore, such cases could lead to higher gene expression noise and yield a more heterogeneous population of cells. It should also be noted that longer residence times are more likely to cause transcriptional bursts (Raser and O’Shea, 2005), which is another notion that contributes to noise in gene expression. On the other hand, shorter DNA residence times are suggested to lower gene expression noise (Azpeitia and Wagner, 2020). Our study sheds light on DNA residence time distributions and may explain the diverse noise levels in gene expression responses.

One of the prominent features of our previous coarsegrained model was the global chromosome organization. Although that system allowed the modeling of the role of nonspecific affinities, selected bead size for TF binding domains led to multiple interactions facilitating the chromosomal collapse. Extreme compaction of the chromosome and resulting residence times of TFs would exceed simulation durations, making it unlikely to obtain RT patterns. By lowering the bead size of TFs, we minimized multiple binding and over-compaction. That also enabled the decoupling of 3D genome organization from RT patterns, increasing the control over other variables.

Similar to any MD study, this work has some limitations. Most prominently, the simplification of the cell for coarse-grained MD simulations removes most of the complexity possessed by the actual cellular systems. Additionally, we used a single type of TF for all MD simulations with discrete DNA binding affinities. In combination, reduced complexity may explain the lack of observation of power-law behavior, which was attributed to continuous TF-DNA affinities in experimental studies. Of course, the cellular complexity is also persistent in the experimental setup. Therefore, other factors, such as fluctuating TF levels and dynamic chromosomal landscape, might be crucial for power-law behavior. However, in our simulations TFs, even with discrete affinities, exhibited multi-exponential behaviors due to emergence, suggesting a power-law pattern is highly possible with some additional complexity even without continuous affinity levels. While this notion does not dismiss the continuum model, it provides an alternative interpretation for the observed power-law behavior.

On the other hand, MD simulations provide distinct advantages. Most prominently, experiments have to rely on approximations and assumptions due to current technological limitations such as indirect measurement techniques. Contrarily, simulations allow direct measurements, providing much higher precision and accuracy. Indeed, the lack of direct measurement necessitates a subsequent correction step, which can introduce biases and lead to inaccurate interpretations of the system or behavior. Correction of the observation data is often not required for MD simulations, resulting in bias-free or less-biased measurements. In fact, we might speculate that the power-law interpretation of residence times distributions in experimental studies might well be due to such biases introduced by corrections rather than the actual underlying TF residence behavior.

In this study, we demonstrate that TFs follow multiexponential patterns with discrete affinities between DNA and TFs in our MD model. We demonstrate how binding affinity and concentration of TFs dictate these behaviors. There are notable implications of this behavior on biological systems such as cellular heterogeneity and noise in gene expression. Isolated from cellular complexity and continuum model, model homodimeric TFs exhibit multi-exponential patterns even with single and binary affinities. This type of behavior might be one of the underlying reasons for gene expression noise and consequent cellular heterogeneity and contribute to cellular differentiation and help overcome evolutionary bottlenecks. Moreover, the distributions of TF-DNA residence times may help explain discrete transcriptional bursts. Lastly, DNA-protein clusters in bacterial chromosomes could drive Topologically Associated Domain-like domain formations and further affect the regulation of gene expression. Overall, our findings on the TF residence time behaviors and their driving factors might contribute to a more comprehensive understanding of gene expression and regulation.

## Methods

### Establishing the system

We used a modified version of our previous coarsegrained model bacterial system mimicking an *Escherichia coli* (*E. coli*) bacterium. The model system includes an adequately relaxed chromosome with uniformly distributed binding sites confined within cell wall-like boundaries resembling that of a rod-shaped bacterium. Generic homodimeric transcription factors with various concentrations (10 *−* 60*μM*) were placed randomly in the volume created by confinement.

### Modelling the DNA polymer

For coarse-grained modeling, we used 10 base pairs (bp) *≈* 1 bead “Kremer-Grest” (KG) model for DNA where 1 bead has a diameter of a *σ* in Lennard-Jones (LJ) units corresponding to *∼* 3.4*nm*. We set the persistent length to 15 beads corresponding to *∼* 50*nm* consistent with a double helix DNA molecule. We established an *N* = 12000 KG bead DNA model where we maximized the number of binding sites and left sufficient spacing for DNA segmental flexibility to minimize the impacts by bridging the binding sites. The initial circular DNA structure was compacted using self-attractive forces to reduce its size to fit within the available volume, similar to our previous work (Koşar et al., 2022). The binding sites are composed of three beads (as opposed to two beads in our previous work), increasing the likelihood of maximum interaction with the TFs. 150 binding sites are placed uniformly along DNA with 80-bead spacing.

### Constructing the confinement

To mimic the rod-like shape of the bacterial cell wall, we first built an open-ended cylinder (or simply a pipe) with a diameter of *R* = 2*r* and a height of 2*R*. Then, two semi-spheres with radiuses of *r* are used as caps to close the endings of the cylinder. The overall structure has a volume of 2*/*3 *× π × R*^3^. The radius (*r*) was set to provide a 1% DNA volume fraction to match that of *E. coli*. Therefore, *r* corresponds to *∼* 30*σ* in LJ units (*∼* 100*nm*) for a DNA polymer length of *N* = 12000. The fixed beads of the cell wall are placed dense enough to provide effective boundaries for DNA and proteins. Nevertheless, there were an insignificant number of TF leaks (*<* 2%) for extended simulations.

### Designing the transcription factors

We designed a generic model of homodimeric TFs in which a TF has two identical binding domains and a hinge domain with no affinity to DNA. Binding domains were placed around the hinge domains with 90^*°*^ angles in a semi-flexible fashion with 12 beads of persistent length. Radii were set to 0.5*σ* and 0.21*σ* for hinge and binding domains, respectively (Figure 1A). We used relatively small binding domains to minimize multiple interactions by single binding domains and prevent over-condensation of the nucleoid. The exact sizes and angles were used to ensure the well-fittings of dimeric TFs into three-bead binding sites. We employed four different concentration levels (i.e.,10*μM*, 20*μM*, 40*μM*, 60*μM*). The corresponding number of TFs for the given concentrations was calculated using the volume provided by the confinement. TFs were distributed into the volume at random coordinates.

### Modelling the TF-DNA affinities

We used a fixed specific interaction potential of *U*_*sp*_ = 4*kT* per bead, energy high enough for robust binding and also low enough to allow unbinding in the time frame of our simulations, enabling us to extract residence time patterns. Varying nonspecific interaction potentials in the range of 1 *−* 4*kT* allowed tracking of residence times at distinct local compaction levels and diverging cluster sizes, as well as differing protein distribution over DNA. For the distributed affinity cases, there was no specific binding potential. Instead, affinities inclusively followed the given ranges of 1 *−* 4*kT* and 2 *−* 5*kT* with the increment of 0.25*kT*, resulting in 13 distinct affinity levels throughout DNA to ensure either Gaussian or uniform distributions.

### Retrieving the residence durations and generating the distributions

Transcription factors are considered bound under the condition that at least one of two binding sites is in direct contact with the DNA polymer (Figure 1B). For each time point, each protein is marked either bound or unbound (Figure 1C). Then, the duration of each uninterrupted bound state was calculated (Figure 1D). Each time a particular residence duration was encountered, the corresponding occurrence was incremented by 1. We then utilized pooled occurrences, containing the number of occurrences for each possible duration (1 *− tmax*), for analyzing residence patterns of transcription factors from simulations with distinct parameters. Obtained data is then minimized to ensure one occurrence value has only one corresponding duration by taking the median of the durations (Figure 1E). This step was necessary for equation fits and visualizations.

### Fitting equations to the residence time distributions

We used several equations to interpret the behavior of transcription factors. Initially, we used single exponential decay *c ×* exp(*−t × k*) and power-law *c × t*^*−k*^ where *c* stands for coefficient, *t* for duration in arbitrary units, *k* for decay rate (i.e., *τ* ^*−*^_1_). Due to sub-optimal fitting with these equations, we included double, triple, quadruple, and pentuple exponential decays in the form of

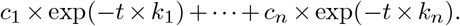

The resulting fits were then graphed and used for calculating the normalized Residual Sum of Squares (RSS). Normalization of RSS values was achieved in two steps. First, occurrence counts are divided by the maximum of occurrence values to normalize occurrences, preventing a higher number of events leading to higher RSS.

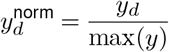

Where 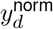 is the normalized occurrence for duration *d, y*_*d*_ is the occurrence value for duration *d*, max(*y*) is the maximum occurrence value in the set of all occurrences.

Then, RSS is divided by the ratio of duration points with non-zero occurrences to prevent more observation points leading to higher total RSS.

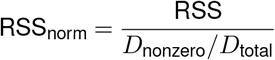

Where, RSS_norm_ is the normalized residual sum of squares, RSS is the residual sum of squares, *D*_total_ is the total number of duration points, *D*_nonzero_ is the number of duration points where occurrences are not zero.

### Clustering the transcription factors

Transcription factors within the threshold distance of each other are accepted to be part of the same cluster. We have used our own deterministic, non-centralized, and vectorized clustering algorithm. We set the threshold distance to 2.1*σ* in LJ units in agreement with visual inspections. For the overview of the system, the minimum number of transcription factors to form a cluster was 12, and for decay analysis, it was 20, ensuring a more accurate estimation of mean lifetimes.

### Cluster analyses

Cluster analyses included size, conformation, surface analysis, and decay rates. The size of a cluster is simply the number of transcription factors within that particular cluster. To classify the conformation (or shape) of a cluster, we first determined three possible formations, namely filamentous, globular, and semi-filamentous. We formed *Rg* tensor for each cluster and evaluated their eigenvalues. To distinguish surface and core cluster proteins, we employed the Convex-Hull algorithm included in the *SciPy* library. Decay rates (reverse of mean lifetimes) for clusters were calculated by fitting a single exponential decay equation to *N* (*t*)*/N*_0_. Decay rates require statistically meaningful numbers for reliable analysis of lifetimes, which led us to select 20 as the threshold for minimum cluster size. We then used linear regression for correlation analysis between cluster dissipation mean lifetimes and their sizes.

## Code Availability

The codes and the processed data necessary to reproduce the analyses and the graphs are available at https://github.com/Zaf4/residence2.

## Acknowledgements

ZK thanks Alper Genceroglu, Irem Niran Çagıl, Leyla Yalçınkaya, Tutku Muratoglu, and Fatma Chafra for their review of the initial manuscript.

## Author Contributions

ZK designed the study, created the models, ran the simulations, performed the data analysis and visualizations, constructed the figures, and wrote the manuscript. AE conceptualized and supervised the project, and revised the manuscript.

