## Supplementary Information for "Protein-DNA Clusters Explain the Non-Exponential DNA Residence Time Distributions of Transcription Factors"

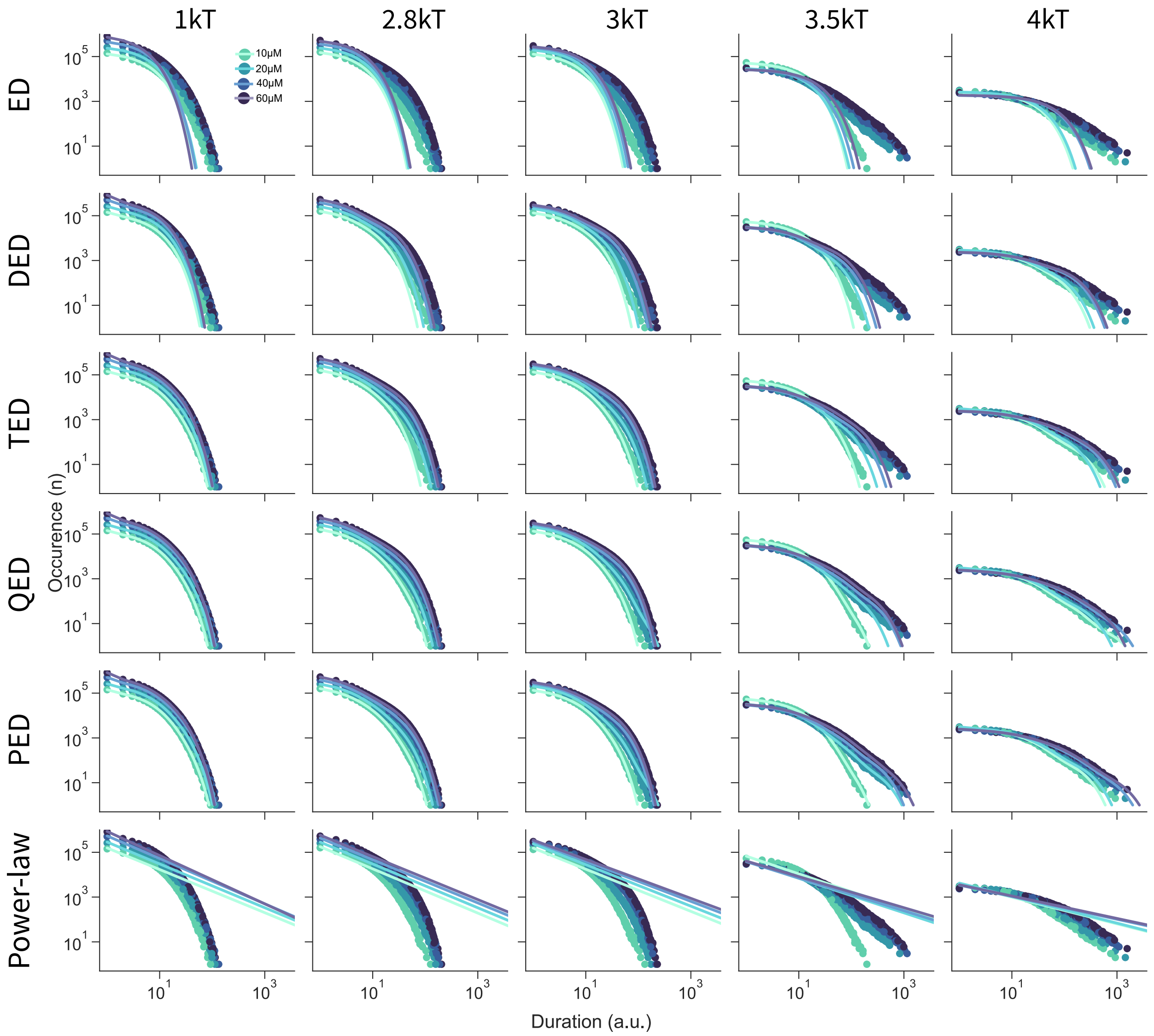


**Figure S1. Residence behaviors of transcription factors**

Distribution of transcription factor residence times in arbitrary unit (a.u.) at various nonspecific binding affinities and fits of Exponential Decay (ED), Double Exponential Decay (DED), Triple Exponential Decay (TED), Quadruple Exponential Decay (QED), Pentuple Exponential Decay (PED), and Power-law equations. Note that specific binding energies are $4kT$ for all the systems.

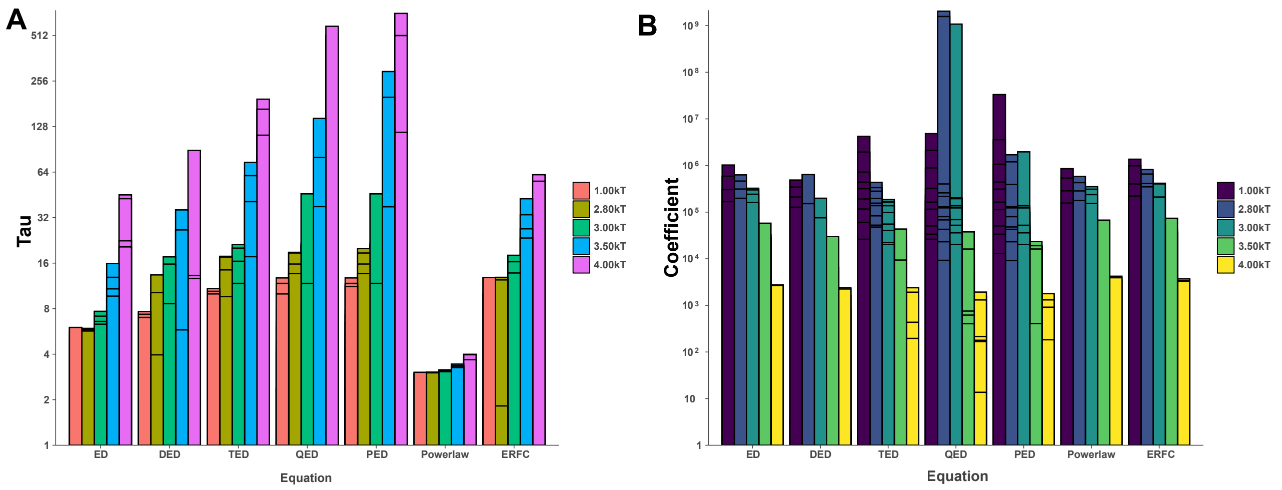


**Figure S2. Exponents of equation fits**

Exponents values of (**A**) Taus and (**B**) Coefficients for each equation with different nonspecific energy levels.

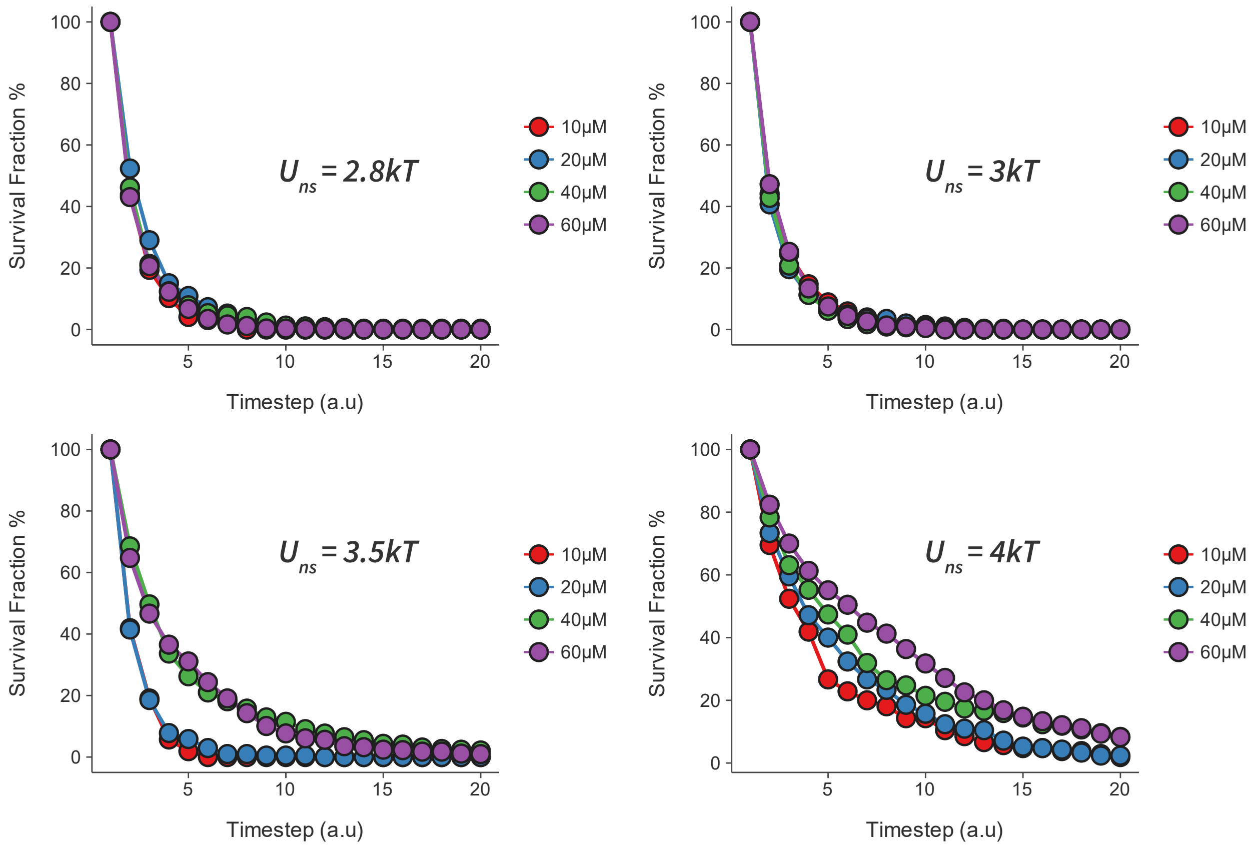


**Figure S3. Survival fractions of transcription factors**

Decay graphs and survival fractions of transcription factors from their binding site on time step 4000 at distinct concentration with affinities with distinct protein-DNA affinity levels. Lines serve to guide the eye.
